# Succession of physiological stages hallmarks the transcriptomic response of fungus *Aspergillus niger* to lignocellulose

**DOI:** 10.1101/806356

**Authors:** Jolanda M. van Munster, Paul Daly, Martin J. Blythe, Roger Ibbett, Matt Kokolski, Sanyasi Gaddipati, Erika Lindquist, Vasanth R. Singan, Kerrie W. Barry, Anna Lipzen, Chew Yee Ngan, Christopher J. Petzold, Leanne Jade G. Chan, Mikko Arvas, Roxane Raulo, Steven T. Pullan, Stéphane Delmas, Igor V. Grigoriev, Gregory A. Tucker, Blake A. Simmons, David B. Archer

## Abstract

**Background:** Understanding how fungi degrade lignocellulose is a cornerstone of improving renewables-based biotechnology, in particular for the production of hydrolytic enzymes. Considerable progress has been made in investigating fungal degradation during time-points where CAZyme expression peaks. However, a robust understanding of the fungal survival strategies over its life time on lignocellulose is thereby missed. Here we aimed to uncover the physiological responses of the biotechnological workhorse and enzyme producer *Aspergilllus niger* over its life time to six substrates important for biofuel production.

**Results:** We analysed the response of *A. niger* to the feedstock *Miscanthus* and compared it with our previous study on wheat straw, alone or in combination with hydrothermal or ionic liquid feedstock pretreatments. Conserved (substrate-independent) metabolic responses as well as those affected by pretreatment and feedstock were identified via multivariate analysis of genome-wide transcriptomics combined with targeted transcript and protein analyses and mapping to a metabolic model. Initial exposure to all substrates increased fatty acid beta-oxidation and lipid metabolism transcripts. In a strain carrying a deletion of the ortholog of the *Aspergillus nidulans* fatty acid beta-oxidation transcriptional regulator *farA*, there was a reduction in expression of selected lignocellulose degradative CAZyme-encoding genes suggesting that beta-oxidation contributes to adaptation to lignocellulose. Mannan degradation expression was wheat straw feedstock-dependent and pectin degradation was higher on the untreated substrates. In the later life stages, known and novel secondary metabolite gene clusters were activated, which are of high interest due to their potential to synthesize bioactive compounds.

**Conclusion:** In this study, which includes the first transcriptional response of Aspergilli to *Miscanthus*, we highlighted that life time as well as substrate composition and structure (via variations in pretreatment and feedstock) influence the fungal responses to lignocellulose. We also demonstrated that the fungal response contains physiological stages that are conserved across substrates and are typically found outside of the conditions with high CAZyme expression, as exemplified by the stages that are dominated by lipid and secondary metabolism.

## Background

Plant biomass is a renewable natural resource that is highly abundant, and is increasingly attracting attention for use of its components in the production of fuels, platform chemicals and materials (1–3). Effective deconstruction of lignocellulose from dedicated feedstock crops such as *Miscanthus*, or agricultural waste such as sugar cane bagasse and wheat straw is becoming increasingly urgent as availability of fossil resources is decreasing (4). Industrial scale use of plant biomass for biotechnologies is hampered by biomass recalcitrance (5,6), whereas nature evolved effective microbial systems to recycle lignocellulose (7–9). To fully exploit such natural deconstruction systems in biotechnology we first need a system-level understanding of microbial actions in response to lignocellulose.

In nature, saprophytic fungi are effective degraders of plant biomass (10–12). Through tightly regulated responses to nutritional and environmental cues they can secrete a large array of hydrolytic enzymes and other effectors (7,13–17). Fungi such as *Trichoderma reesei* and *Aspergillus niger* are widely used in commercial production of these enzymes, due to their high titer secretion of degradative enzymes under industrial conditions (18,19). In addition *A. niger* has a firmly established role as industrial workhorse in the production of organic acids and other high value compounds (19). It is therefore imperative to increase the understanding of the saprotrophic biology of these particular fungi.

In biotechnology, strategies to overcome substrate recalcitrance include engineering of enzyme cocktails and microbial strains for improved degradation as well as improving biomass digestibility via engineering of its composition or structure, or via chemical or physical pretreatment (6,20–22). Pretreatments, currently essential for effective lignocellulose processing, increase biomass digestibility via various mechanisms. Hydrothermal treatments remove hemicellulose components and break linkages between lignin and polysaccharides (23,24) while ionic liquids solubilise wall components and generate, after recovery, insoluble but highly digestible polymers (3,25). However, even small differences in composition and structure of plant cell walls can significantly affect their digestibility (20,26). As ascomycetes have been described to strongly respond to differences in polysaccharides used as carbon source (27–29) and also exhibit partially conserved (substrate independent) responses to complex lignocellulose (30–33), here we investigated the effect of both hydrothermal and ionic liquid pretreated biomass as well as untreated feedstocks on the system-level response of *A. niger*.

The production of plant cell wall hydrolytic enzymes by *A. niger* is a highly orchestrated process. Initial responses of the fungi to biomass include alleviation of carbon catabolite repression and production of a set of scouting enzymes that scavenges for carbon sources (32,34–38). These then mediate release of small soluble carbohydrates from biomass, thereby inducing expression of an appropriate broad set of degradative enzymes via a network of transcriptional regulators (7,14,39–42). Subsequent degradation of lignocellulose components can be expected to reflect the evolved fungal biochemical capacity (27,43). Fungal production of CAZymes and signalling responses to different sources of complex carbohydrates have been documented extensively, often capturing a snapshot of events at one or a selected few moments, such as when the CAZyme expression was peaking, in the cultivation time (28–34,43–46). Knowledge on the wider, and potentially time-staged, physiological responses and degradative strategies of *A. niger* is however still very limited.

To overcome this limitation, here we aimed to generate a systems level understanding of physiological responses of *A. niger* to lignocellulose. Previously, we found that the expression of *A. niger* CAZymes was determined by compositional changes in wheat straw generated by hydrothermal or ionic liquid pretreatments. We demonstrated how a high-resolution time series analysis captured a comprehensive view of the production of CAZymes in response to substrate compositional changes in wheat straw but did not examine other physiological responses (31). Here we build on this approach to elucidate the effect of feedstock identity as well as pretreatment and cultivation time on the wider *A. niger* transcriptional landscape. We applied the high resolution time-staged transcriptome approach to cultivations with *Miscanthus* lignocellulose, both untreated and pretreated with ionic liquid or hydrothermally. Using this first dataset of the *Aspergillus* transcriptome on *Miscanthus* we furthermore aimed to uncover conserved fungal cell biological responses that are important to biomass degradation. The physiological responses to lignocellulose identified here will improve understanding of fungal degradative strategies, and potentially aid understanding on the requirements for development of *Aspergillus* strains for bioprocessing.

## Results and discussion

### *A. niger* cultivation over time is affected by pretreatment of *Miscanthus* that influences its composition and digestibility

*A. niger* was exposed to knife-milled *Miscanthus* (KMM) as well as ionic liquid (IL) and hydrothermally (HT) pretreated *Miscanthus* for up to 5 days to identify the system-level fungal responses upon exposure to these substrates (Fig. 1a). Time-courses were terminated earlier where RNA of sufficient quantity and quality could no longer be obtained, i.e. after 24 h exposure of mycelium to HT pretreated *Miscanthus* (HTM) or 2 days to KMM, which suggested reduced fungal viability under these conditions.

**Fig. 1.**
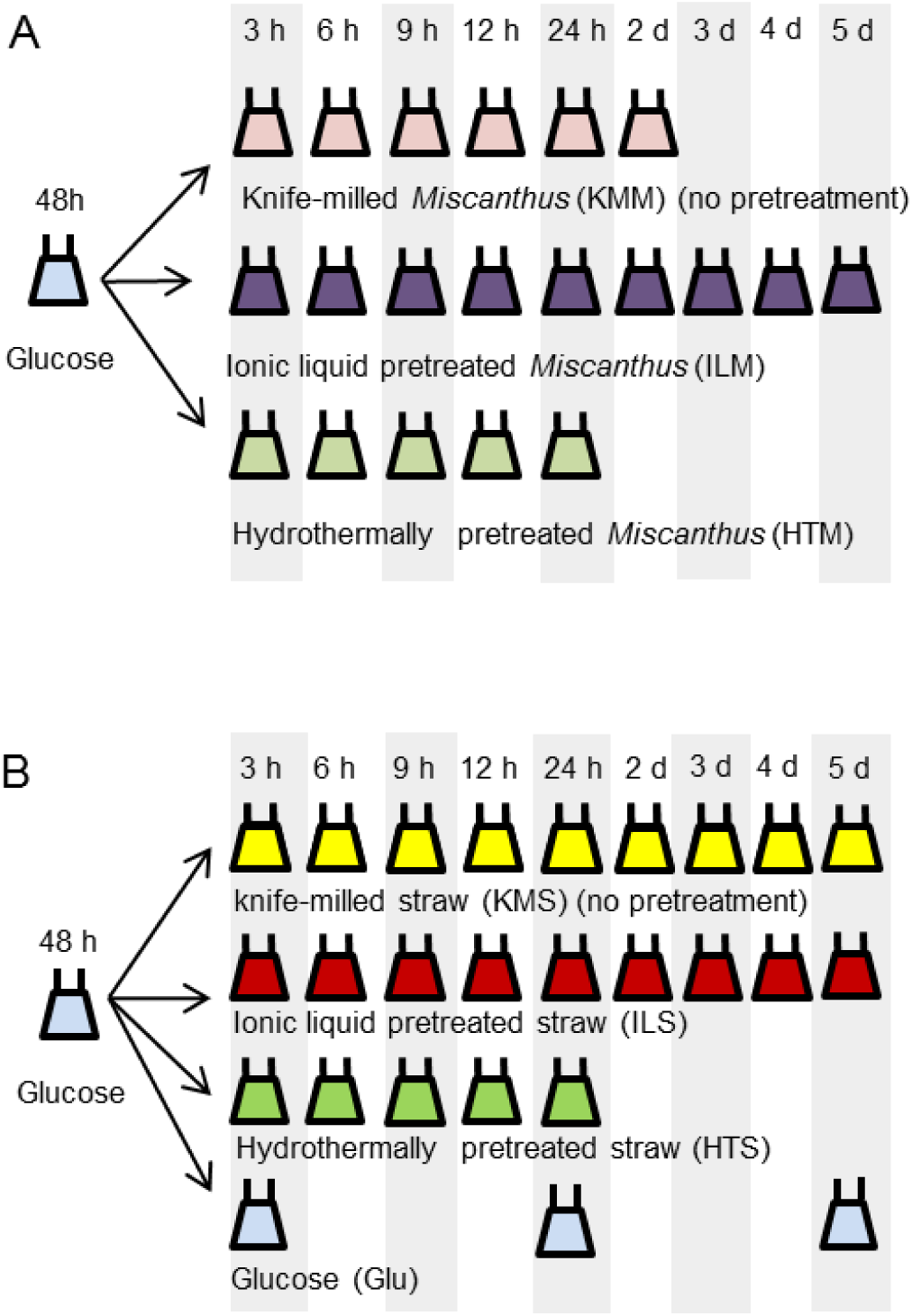
Schematic representation of experimental setup. **a** *A. niger* was cultivated in shake flasks with glucose as carbon source, after which mycelium was transferred to shake flask containing either knife milled *Miscanthus* (KMM), or KMM pretreated with ionic liquid (ILM) or hydrothermal treatment (HTM). Mycelium was harvested at indicated time points after which RNA was isolated and sequenced. This dataset was compared to data obtained with a similar experiment **b** using knife milled wheat straw (KMS) and pretreated straw (ILS, HTS) (31). Panel b is reproduced from (31) under a CC BY ((https://creativecommons.org/licenses/by/4.0/) license.

HT pretreatment of *Miscanthus* reduced the arabinose and galactose content ≥10-fold and the xylose content 2-fold (Fig. S1a), indicating loss of hemicellulose and pectin, while IL pretreatment reduced cellulose crystallinity (Fig. S1b). As expected both pretreatments improved substrate saccharification with a standard commercial enzyme cocktail (Fig. S1c). Fungal degradation of HTM appeared to be less than that achieved with the commercial enzyme saccharification. The xylan detected in the solid fraction retrieved from cultures of *A. niger* with KMM reduced by 26 % over the first 24 h of cultivation, whereas in HTM it reduced by only 18 %, suggesting less degradation of HTM than KMM during cultivations (Fig. S1d). In contrast, xylan content of ILM decreased by 71% after 24 h cultivation, indicating that its increased digestibility in enzymatic saccharification also occurred in the fungal cultivations. Levels of monosaccharides in culture filtrates support increased ILM digestibility in fungal cultivations compared to KMM. In the 9-24 h time points of exposure to ILM, up to 2.2 mM of xylose and arabinose was detected (Fig. S1e), whereas monosaccharide concentrations in other conditions were <100 mM. This suggests high sugar release via effective ILM digestion and accumulation to levels reported to be high enough (47) to activate carbon catabolite repression.

The overall effect of HT and IL pretreatment on *Miscanthus* feedstock composition and digestibility, and its subsequent degradation in fungal cultures, was similar to our wheat straw study (31). However, feedstock identity affected the end result of pretreatments and subsequent fungal cultivation; *Miscanthus* was less digestible than knife milled wheat straw (KMS) and pretreatments only partly remedied this. For example, saccharification with commercial enzymes released 4-fold less sugar from KMM than from KMS but 2-fold less from HTM than from HT pretreated straw (HTS), with no difference for ILM and IL pretreated straw (ILS).

### Overview of gene expression during cultivation on complex substrates

We investigated the transcriptional landscape of niger to untreated and pretreated Miscanthus to elucidate the effect of feedstock and pretreatment on gene expression, and to identify the system-level fungal responses to cultivation. We analysed the data together with that of cultivations on untreated and pretreated wheat straw from our previous study (31), where the broader physiological responses beyond CAZymes had not yet been analysed. This resulted in a data matrix of 47 conditions of *A. niger* cultured with a particular (pre- or untreated) substrate (Fig 1). Hierarchical clustering of the transcript levels of all genes (Fig. 2a), showed that pretreatment had the most discriminatory effect, as demonstrated by clustering of selected IL pretreated conditions with glucose-grown conditions (Cluster 1, C1, indicated in blue). The second most influential contribution to differences between samples was cultivation time; early (3-6 h) time points clustered together, as do 9-12 h conditions, and together these group (C3, red) separately from later time points (C2, green). Surprisingly, lignocellulose feedstock identity (i.e. *Miscanthus* or wheat straw) had an overall minor effect on clustering pattern. Subsequent PCA confirmed separation of conditions on the first to third principal component (Fig. 2b,c) corresponded mainly to effects of pretreatment and cultivation time, with follow-on detailed analysis revealing feedstock effects.

**Fig. 2.**
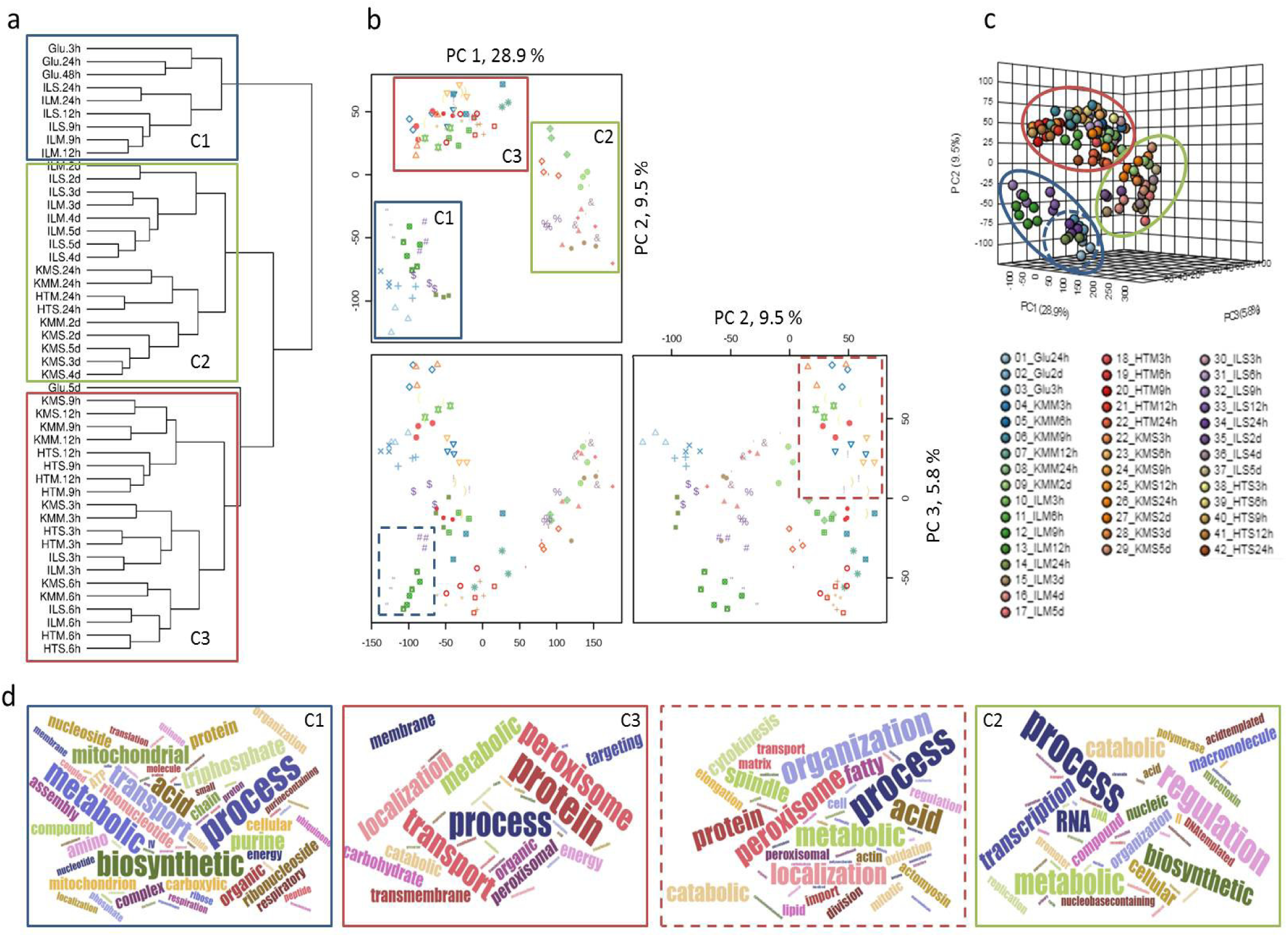
Hierarchical clustering and PCA analysis illustrate time, pretreatment and substrate dependant clustering patterns. With **a** hierarchical clustering of all conditions using the log transformed and quartile normalised mean FPKM values that were ^3^ 1 in any of the time points on any media, **b** and **c** results from PCA showing separation in distinct clusters over the first 3 components in 2D and 3D respectively. Ellipses indicate 95% confidence intervals. Clusters that are discussed in the main text are labelled C1-C3 and indicated with coloured panels, with dashed lines representing sub-clusters. Panel **d** summarises relative abundance of GO-terms identified as linked to the clusters via word clouds.

The genes and processes underpinning the observed variation in gene expression between sample groups were identified via analysis of the PCA loadings, (which identifies genes whose expression levels contributed highest to the sample group distribution) via gene transcription patterns and GO-term enrichments. Samples from cultures grown on glucose and those cultivated on IL pretreated substrates for 9 h, 12 h and 24 h, cluster together (C1) on PCA1 and PCA2. Genes with highest loadings for this group, whose expression levels contributed highest to the clustering, were enriched for those annotated with GO-terms for respiration, e.g. mitochondrial function including transport, organisation, ATP metabolism, as well as biosynthesis of compounds as amino acids and organic acids. IL 9 h and 12 h time points distinguished themselves (Fig. 2b,c) from the glucose grown samples via enrichment of GO-terms associated with translation. Identified GO-terms are summarised in word clouds (Fig 2d).

The 3 h to 12 h time points on lignocellulose (except ILM and ILS 9-12 h) formed cluster C3. Genes with the highest loadings were enriched for those annotated with GO-terms for carbohydrate metabolism, cellular respiration, protein transport and peroxisome function or organisation. Closer inspection revealed that 3 h and 6 h time points on lignocellulose formed a subcluster C3a (Fig 2b, indicated in dashed red), which largely consisted of the genes with the highest loading enriched for those annotated with GO-terms for fatty acid beta-oxidation, peroxisome organisation, cell division and cytoskeleton (re)organisation.

Lignocellulose samples at 24 h or later (except ILM and ILS 24 h) clustered together (Cluster C2). Genes with the highest (≥0.1) loadings were enriched for those annotated with GO-terms for catabolism of polysaccharides, aromatic amino acids and other aromatic compounds, the biosynthesis of secondary metabolites as terpenoids and mycotoxins, modification of fungal cell walls, chromosome organisation and regulation of metabolic processes. This coincides with enrichment of oxidoreductase and glycoside hydrolase activity associated terms.

### Initial response to lignocellulose involved increased fungal lipid metabolism

The initial transcriptional response to lignocellulose was conserved over feedstocks and pretreatment, as demonstrated by clustering of the early (3-6 h) time points in PCA and hierarchical clustering. For genes with reduced transcript levels, GO terms for RNA helicase activity were enriched at the 3h time-point (Table S1), as well as those for RNA processing - specifically processing and maturation of non-coding and ribosomal rRNA - and ribosomal subunit biogenesis, assembly and localisation, together suggesting a conserved reduction in overall translational activity upon transfer to lignocellulose. For genes with increased transcript levels, enriched GO-terms (Table S2) strongly represented lipid catabolism, specifically fatty acid beta-oxidation and catabolism of the short chain fatty acid acetate (Fig. 3a). This is in accordance with the results from the PCA. In addition, the GO-term for branched-chain amino acid breakdown was enriched at the 3 h time points, which could be caused by breakdown products of valine and isoleucine which are processed via fatty acid beta-oxidation (48). Metabolic mapping highlighted significant enrichment of the butanoate metabolic pathway, a short chain fatty acid, in the early time points, as well as gluconeogenesis pathway enrichment (Table S3).

**Fig. 3.**
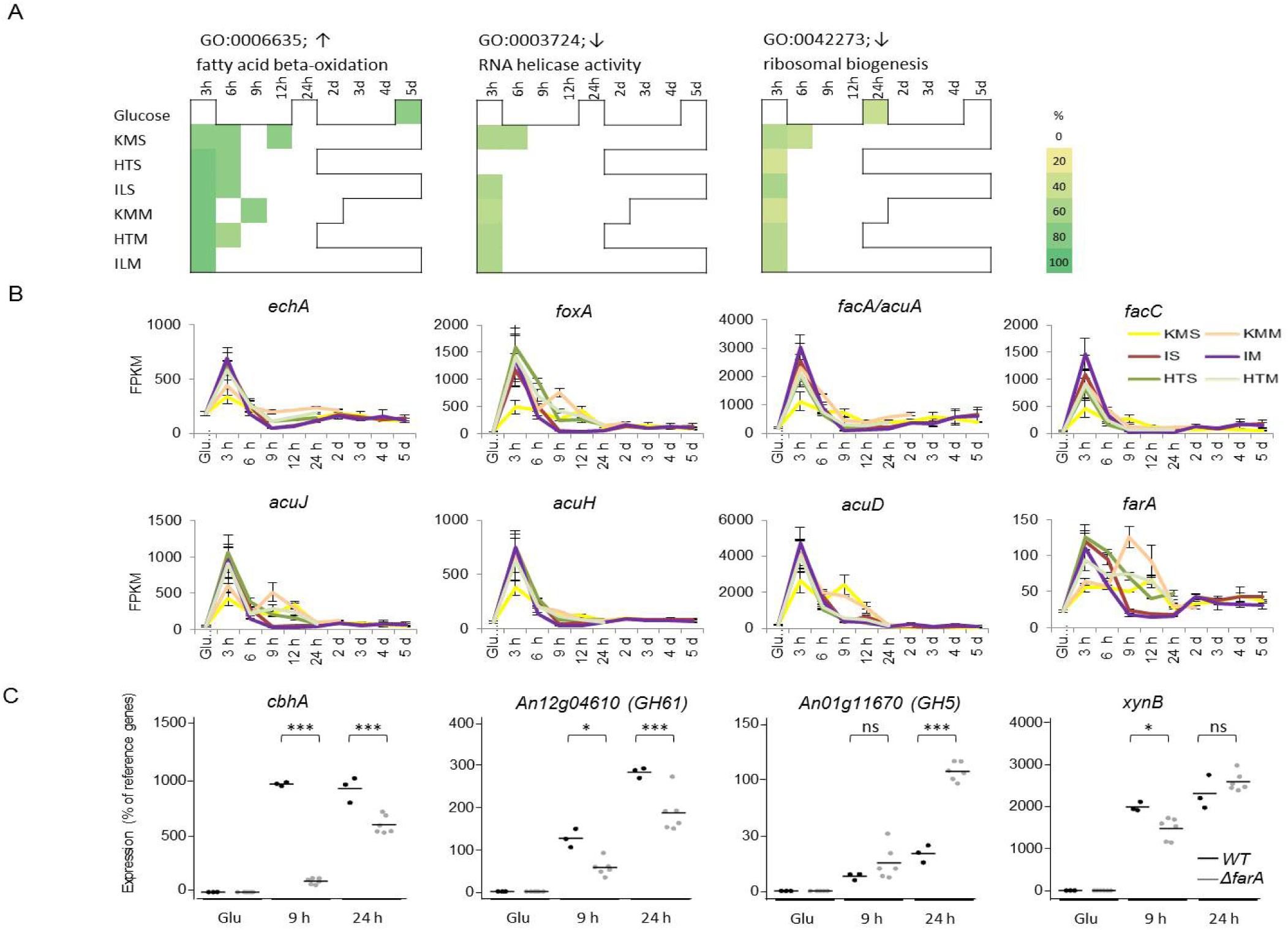
Lipid metabolism as general response in early time points. With **a** enrichment of selected GO-terms in early time points, **b** transcript levels of genes involved in fatty acid β-oxidation and acetate metabolism, as detected by RNASeq, **c** *cbhA, GH61a* (*An12g04610*), *GH5 (An01g11670)* and *xynB* transcript levels in the wild-type (black) and *farA* (grey) deletion strain grown on glucose or wheat straw (9 and 24 h) as detected by qRT-PCR, mean indicated by bar, results from statistical testing is indicated by ns (*p*>0.05), * (*p*<0.05), ** (*p*<0.01) or *** (*p*<0.001).

Fatty acid metabolism has been investigated most thoroughly in *A. nidulans*, and many orthologous genes are present in *A. niger*. Short chain fatty acids are oxidized to produce acetyl-CoA in the mitochondrion by, among others, ScdA and EchA, while long chain fatty acids are oxidised in the peroxisome by FaaB and FoxA (48,49). Transcript levels of corresponding *A. niger* genes (Fig. 3b) showed a peak at 3 h, indicating that both short-chain and long-chain fatty acid beta-oxidation is activated. Acetyl-CoA is metabolised via the tricarboxylic acid (TCA) cycle in the mitochondrion. TCA cycle intermediates are used for gluconeogenesis, and are replenished in the peroxisome by AcuD and AcuE. To enable acetyl-CoA shuttling between cell compartments, carnitine acetyltransferases FacC, AcuJ (50,51) interconvert acetyl-CoA and acetylcarnitine, which can cross membranes using transporters such as AcuH. The *A. niger* genes encoding these proteins, as well as An11g02590.Aspni5_55954, encoding PexK required for peroxisome biogenesis during fatty acid β-oxidation (52,53), all show a similar expression profile with a peak at 3 h (Fig. 3b). Genes specific for acetate metabolism, *facC* (54), and *acuA* (ortholog of *A. nidulans facA)*, also show a peak in transcript levels at the 3 h (Fig. 3b) indicating that the acetate metabolism pathway is specifically activated during early exposure to complex lignocellulose substrates.

Taken together these data strongly suggest that directly upon exposure to lignocellulose, fatty acid beta-oxidation and subsequent metabolism is increased. During the carbon-starved conditions experienced at that point, this could provide the fungus with a source of energy and result in the generation of glucose via gluconeogenesis. The response is conserved over all substrates analysed here, indicating a universal process consistent with the carbon starvation experience. In line with this, enrichment of almost all of these GO-terms was also observed in the 5 d glucose culture where glucose is depleted and increased transcription of genes linked to fatty acid metabolism in response to carbon starvation has been reported for Aspergilli (16,55) as well as in other fungi (56,57). A role for carbon starvation in induction of plant polysaccharide degradative enzymes was previously established in *A. niger* (34,38), *A. nidulans* (36,37,58) and *N. crassa* (28,59), which involved low-level upregulation of a subset of genes encoding CAZymes active on plant polysaccharides, the ‘scouting enzymes’. Besides generating energy for cell maintenance, fatty acid metabolism may generate energy required for production of these scouting enzymes or assist in establishing fungal colonisation of lignocellulose. To test this hypothesis, we analysed the role of fatty acid metabolism in hydrolase production via deletion mutants of the ortholog of the *A. nidulans* gene *farA*, encoding the inducible regulatory activator responsive to both short- and long-chain fatty acids FarA (60).

### The ortholog of *A. nidulans* FarA contributes to timely establishment of hydrolytic capacity

The expression of *farA* strongly increased at 3 and 6 h on all lignocellulose substrates, and in addition at 9 h for KMM (Fig. 3b). As this gene is the ortholog of the confirmed regulatory activator of fatty acid metabolism in *A. nidulans*, it was an ideal candidate to assess an influence of fatty acid metabolism in establishing hydrolytic capacity via *farA* deletion.

Gene expression in two independently generated *farA* deletion mutants and its wild-type parent strain were analysed by qRT-PCR after exposure to lignocellulose. After 9 h, the *farA* deletion strains had significantly (*p*<0.05) lower transcript levels for the cellulolytic enzyme encoding genes *cbhA* and *GH61a* (*An12g04610*): compared to the wild-type, mean levels of transcripts were 10-fold and 2-fold less induced respectively (Fig. 3c). After 24 h transcription levels of these genes were more similar with <2-fold difference. No pronounced differences (*p*>0.05 and/or <2-fold change) were found for transcription of *xynA* or *xynB* at 9 h or 24 h, *w*hile transcript levels of GH5 *An01g11670* were similar in both strains at 9 h but were increased 3-fold in expression in the *farA* deletion strain after 24 h. These results indicate that the initial substrate-independent lipid metabolism via FarA affects the subsequent induction of a subset of genes encoding CAZymes during exposure to lignocellulose. The results also suggest the mechanism may not be via a direct effect on scouting enzyme transcript levels in early time points, as no difference for initial transcription levels of a GH5 scouting enzyme was detected at 9 h. An indirect effect would be consistent with the lack of FarA binding sites (CCTCGG (60)) in the 1 kbp upstream promoter regions of the *cbhA* and *GH61a* genes. Rather, the effect could be through translational effects, or cumulative effects that eventually result in a delayed fungal lignocellulose colonisation and thus delayed induction of genes that encode enzymes routinely observed only under full induction on this substrate (34,38), such as *cbhA* and *GH61a*.

### Pretreatment and feedstock effects dominate gene expression in middle-stage cultures on lignocellulose

Our initial multivariate analysis indicated that the 9-12 h time points on all substrates represent established (middle stage) cultures on lignocellulose with the most active carbohydrate degradation and metabolism, cellular respiration and protein transport (Fig. 2). PCA analysis of gene expression levels indicated at least 27% of variation in gene expression in these time points resulted from differences between IL versus other cultures, with ≥ 9% of variation explained by HT versus other cultures and ≥ 5% by feedstock (Fig. 4a).

**Fig. 4.**
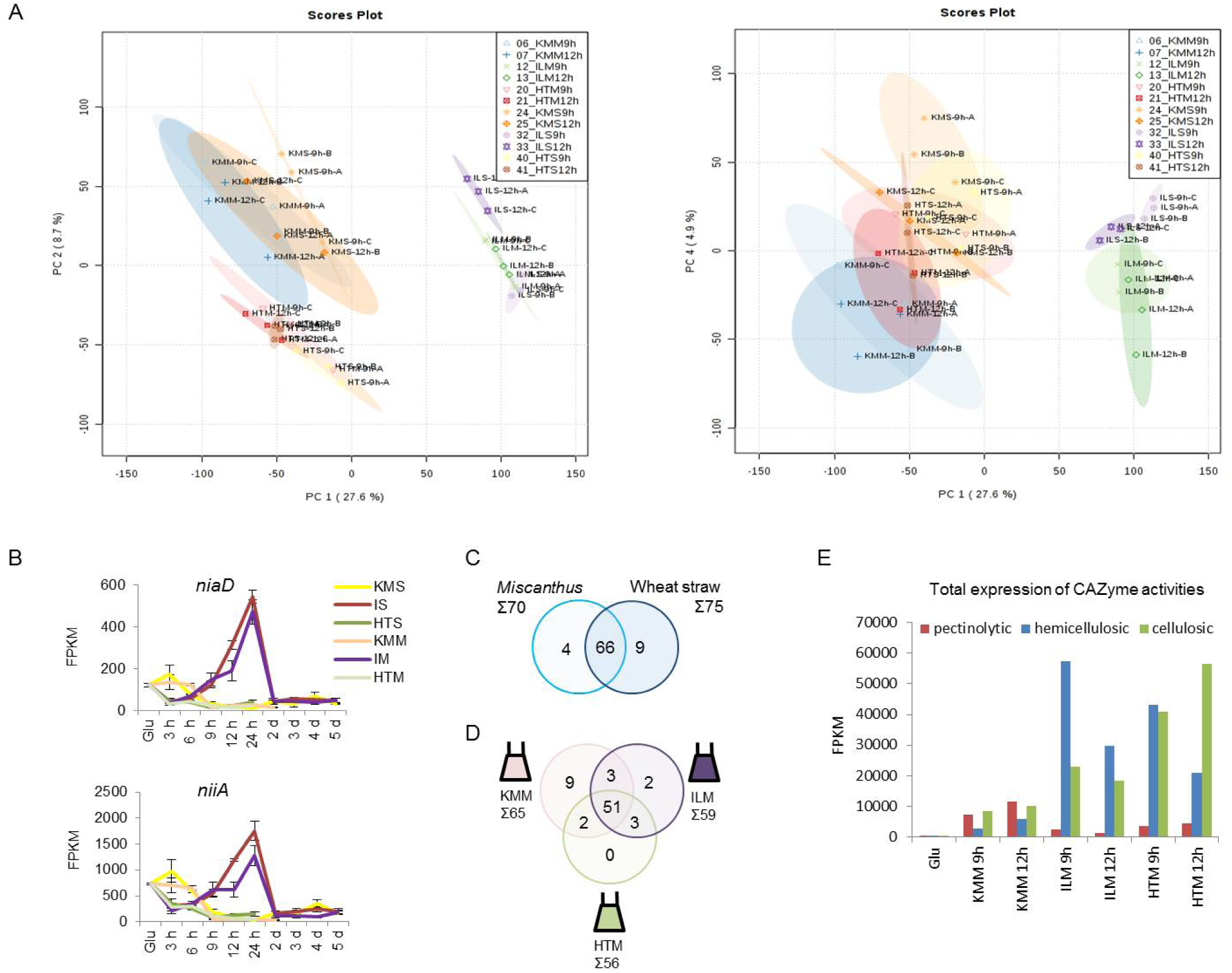
Feedstock and pretreatment effects in established cultures on *Miscanthus* include nitrate assimilation and CAZyme expression. **a** PCA analysis of variation in gene expression in 9h and 12h *Miscanthus* and straw cultivations, demonstrating pretreatment (PC1, PC2) and feedstock (PC4) effects. Ellipses indicate 95% confidence intervals. **b** Expression of nitrate assimilation genes *niaD* and *niiA* identified via RNA-Seq, values are mean ± st. error, **c** Venn diagram illustrating overlaps in total number of induced plant polysaccharide-active CAZyme encoding genes during exposure to *Miscanthus* and wheat straw, **d** and between induced genes on untreated and pretreated *Miscanthus* substrates, **e** total expression, as sum of FPKM, for genes encoding hydrolytic enzyme activities over the 9h and 12h *Miscanthus* time courses. See Table S5 for underlying data.

Genes with positive loadings in the PCA first component that distinguishes IL pretreatment from the other conditions, revealed high expression of a cluster of genes encoding proteins responsible for nitrate uptake and intracellular conversion to ammonium, including nitrate reductase NiaD and nitrite reductase NiiA (61). High expression of this gene cluster was shared with the glucose cultures and with early KMS time points (Fig. 4b). When using nitrogen assimilation activity as a proxy for growth (62), this indicated fungal growth on IL pretreated substrates from 3 h to 24 h, reflecting high digestibility of the IL pretreated substrates (Fig. S1). Genes associated with HT pretreatment, with negative loadings in the second principle component, were enriched in GO-terms for phenylalanine catabolism and oxidoreductase activity.

Genes linked to complex carbohydrate degradation had high loadings on the first principle component (IL pretreatment) and the fourth principle component, separating *Miscanthus* and wheat straw feedstocks. We therefore compared expression of genes encoding plant-polysaccharide degrading CAZymes that showed at least in one of the time-points a high-level induction compared to the glucose preculture (i.e. with a DESeq *p*_*adj*_ < 0.05, FPKM of ≥ 50 on lignocellulose and log2 FC of ≥ 3) (Fig. 4c). The sum of transcript levels of genes encoding plant CAZymes formed a dome-shaped curve during exposure to lignocellulose (Fig. S2b), with a dip in expression levels during the 12-24 h time points in ILM cultures similar as reported for ILS cultures (31), as high IL-pretreated substrate digestibility resulted in accumulation of xylose to levels responsible (47) for carbon catabolite repression (Fig S1). While the overall number of induced CAZyme-encoding genes was very similar on wheat straw and *Miscanthus*, the total expression level on KMM was reduced compared to KMS (Fig. S2).

Over the time-course on *Miscanthus*, 70 CAZyme-encoding genes were induced (Fig. 4c, Table S5). A core set of 51 genes encoding CAZymes with (putative) plant-polysaccharide degrading activity were induced both in cultures grown with untreated and pretreated *Miscanthus* (Fig. 4d). We detected nine genes that were specifically induced on KMM, including six pectinolytic genes. ILM specific induced genes included the α-galactosidase encoding *aglB*, while the α-xylosidase encoding *axlA* was ILM and HTM specific.

We compared the expression levels of the plant encoding CAZy genes induced at the 9 h and 12 h time points as these established cultures showed minimal transcriptional repression by CCR and overall high CAZyme expression levels. Considerably higher average expression levels were found after exposure to ILM and/or HTM when compared to KMM, with induction between 5 to 80-fold (Table S5). Investigating the induced activities (Fig. 4e), a broad set of genes encoding hemicellulosic and cellulosic enzymes had much higher transcript abundance on pretreated substrates when compared to KMM, with ILM displaying the highest transcript abundance of genes encoding hemicellulosic enzymes and HTM the highest transcript abundance of genes encoding cellulosic enzymes. KMM on the other hand displayed the overall highest transcript abundance of genes encoding pectinolytic enzymes, in line with the number of pectinase-encoding genes expressed in this condition. Confirmation that these transcript abundance differences also translated into relative protein abundance and thus degradative capacity was obtained by targeted proteomics. Relative abundance of xylanolytic enzymes XynA and XynB, but not XlnD was found to be higher in culture filtrates of ILM and HTM compared to KMM (Fig. S3). Cellulolytic CbhA, EglA and an AA9 putative LPMO, but not CbhB and EglC, were more abundant in filtrates of HTM than of KMM cultures. Pectinase PelA was most abundant in filtrates from KMM cultures (Fig. S3).

The relatively abundant pectinase production on untreated *Miscanthus* was also observed on wheat straw (KMS) (31) and may be related to the reduction of pectin content by the pretreatments, and resultant lack of inducers for pectinase encoding genes (41). HT-dependent higher production of cellulases could be explained by the relatively higher cellulose content of this substrate and its increased digestibility compared to KMM.

### Feedstock-specific enrichment for mannan degradation

Comparing CAZy gene expression over the time course of cultivations on both wheat straw and *Miscanthus* feedstocks (Fig. 4c), mannan degradation was found to be feedstock-specific. Enrichment of its GO-term (GO:0046355) was specific to the untreated and pretreated wheat straw cultures. Four of five genes annotated with the GO term for mannan degradation were responsible for this enrichment (Table S2). When looking at the expression profile of these genes in detail, induction of transcription of β-mannosidase *mndA* was straw specific, while endo-β-mannanase encoding *man5A* had much higher transcript levels on wheat straw, especially KMS and HTS, compared to *Miscanthus* (Fig. 5a). Targeted proteomics showed a similar trend for the abundance of this protein in the culture filtrate (Fig. 5b) and endo-mannanase activity was detected at higher levels in wheat straw culture filtrates compared to the *Miscanthus* culture filtrates (Fig. 5c). These results confirm the up-regulation of endo-β-mannanase activity is feedstock-specific. Three out of four of the genes responsible for enrichment of the GO term are inducible by mannose or mannan (41). Mannan content of wheat straw and *Miscanthus* is very low (63) and perhaps instead increased accessibility of the mannan in the wheat straw could be a contributory factor to the observed distinct activation of mannan degradation.

**Fig. 5.**
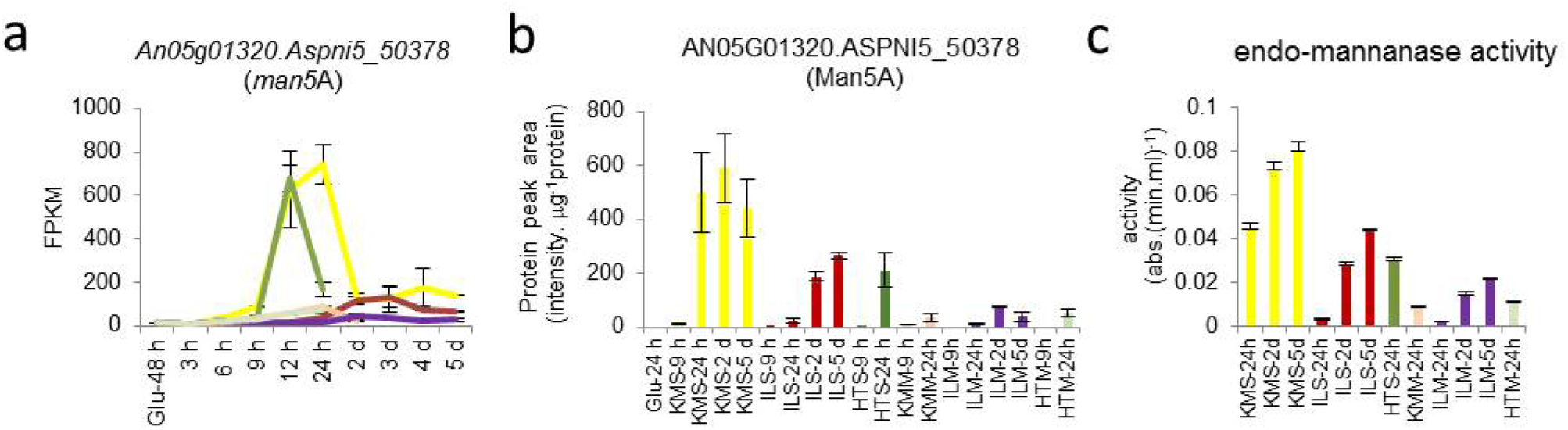
Mannan degradation is feedstock specific. **a** gene expression levels of the endo-β-mannanase encoding gene An05g01320. Aspni5_50378, **b** protein levels of the endo-β-mannanase in the culture filtrate and **c** endo-mannanase activity detected on AZCL-galactomannan. Error bars represent standard errors.

### Transition to starvation and sporulation

The high-level transcription of plant biomass degrading CAZy genes was not sustained across the time-course. A feedstock independent transition to reduced enzyme expression and activation of starvation, sporulation and secondary metabolism occurred between 12-24 h (KM and HT) or 24 h-2 d (IL), as indicated by hierarchical clustering and PCA (Fig. 2).

The transcriptomic data indicated limited growth and primary metabolic activity at the later time points. Expression of nitrogen assimilation related genes *niiA* and *niaD* was limited (Fig. 4b) and expression of the genes encoding key biosynthesis enzymes for the fungal cell membrane component ergosterol (*erg2, erg3, erg11* and *erg25)* was considerably reduced (Fig. 6a). In agreement, steroid biosynthesis (KEGG rn00100) was enriched in the metabolic mapping in pre-, but not post-24h time points. As ergosterol is a marker for fungal biomass (64) the depletion of transcripts suggests limited fungal growth after the transition point. During late (≥ 24 h) time points GO-terms suggested a low metabolic activity: terms enriched in genes with reduced transcript levels included aerobic respiration, specifically citric acid (TCA) cycle and acetyl coenzyme A degradation, oxidative phosphorylation and ATP synthesis (Fig. 6b).

**Fig. 6.**
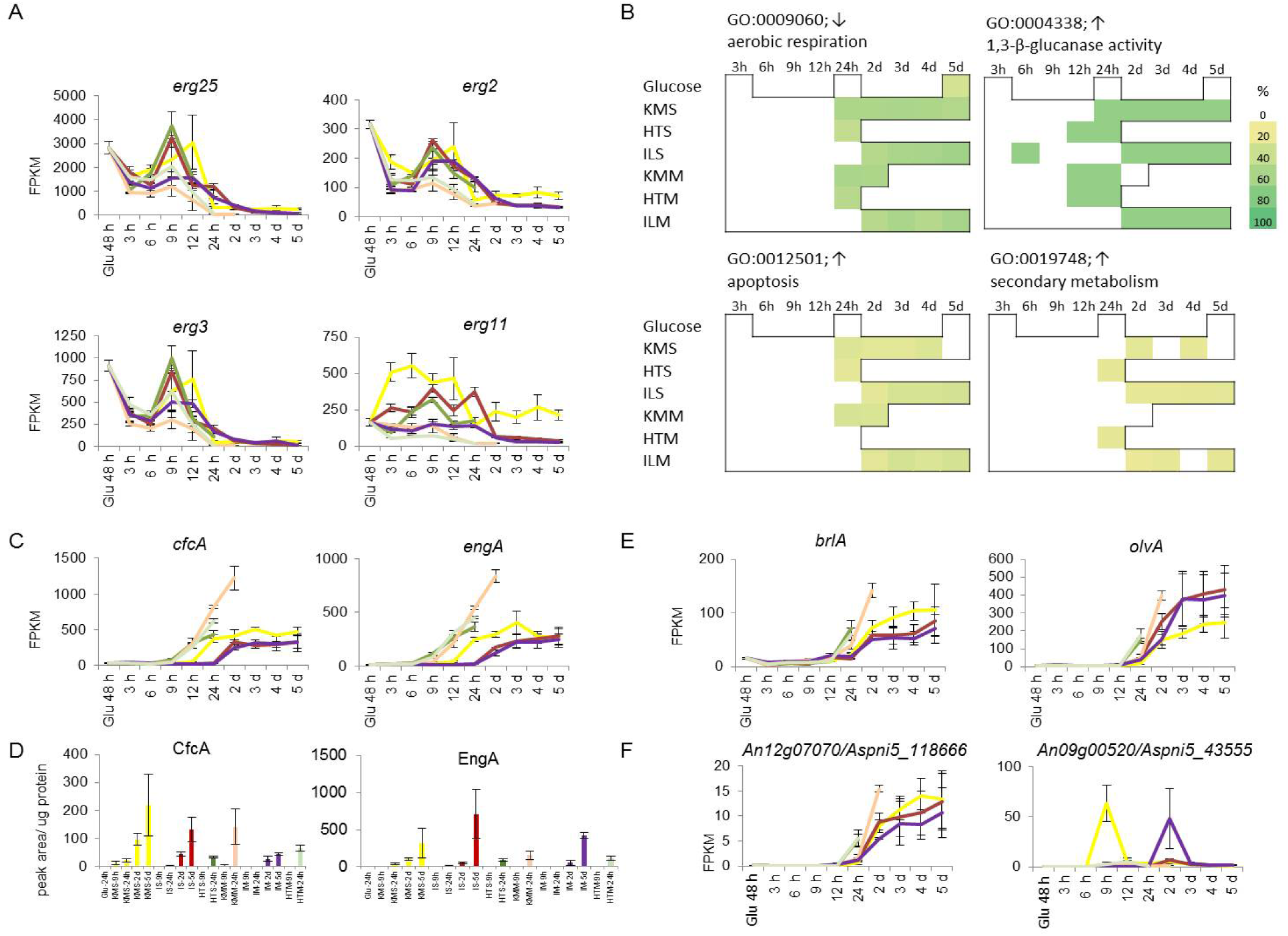
Late time point cultures are starving and induce secondary metabolism. **a** transcript levels of genes encoding ergosterol biosynthesis enzymes, **b** enrichment of selected GO-term in late time points, **c** transcript levels of genes encoding starvation-induced fungal cell wall acting hydrolases, **d** protein level of corresponding fungal cell wall acting hydrolases in the culture filtrates and **e** transcript levels of genes encoding the putative PKS of a sporulation specific secondary metabolite cluster (left), and the NRPS of the gliotoxin cluster (right), with 2 and 7 respectively of the other cluster genes sharing this profile. Error bars represent standard errors.

**Fig. 7.**
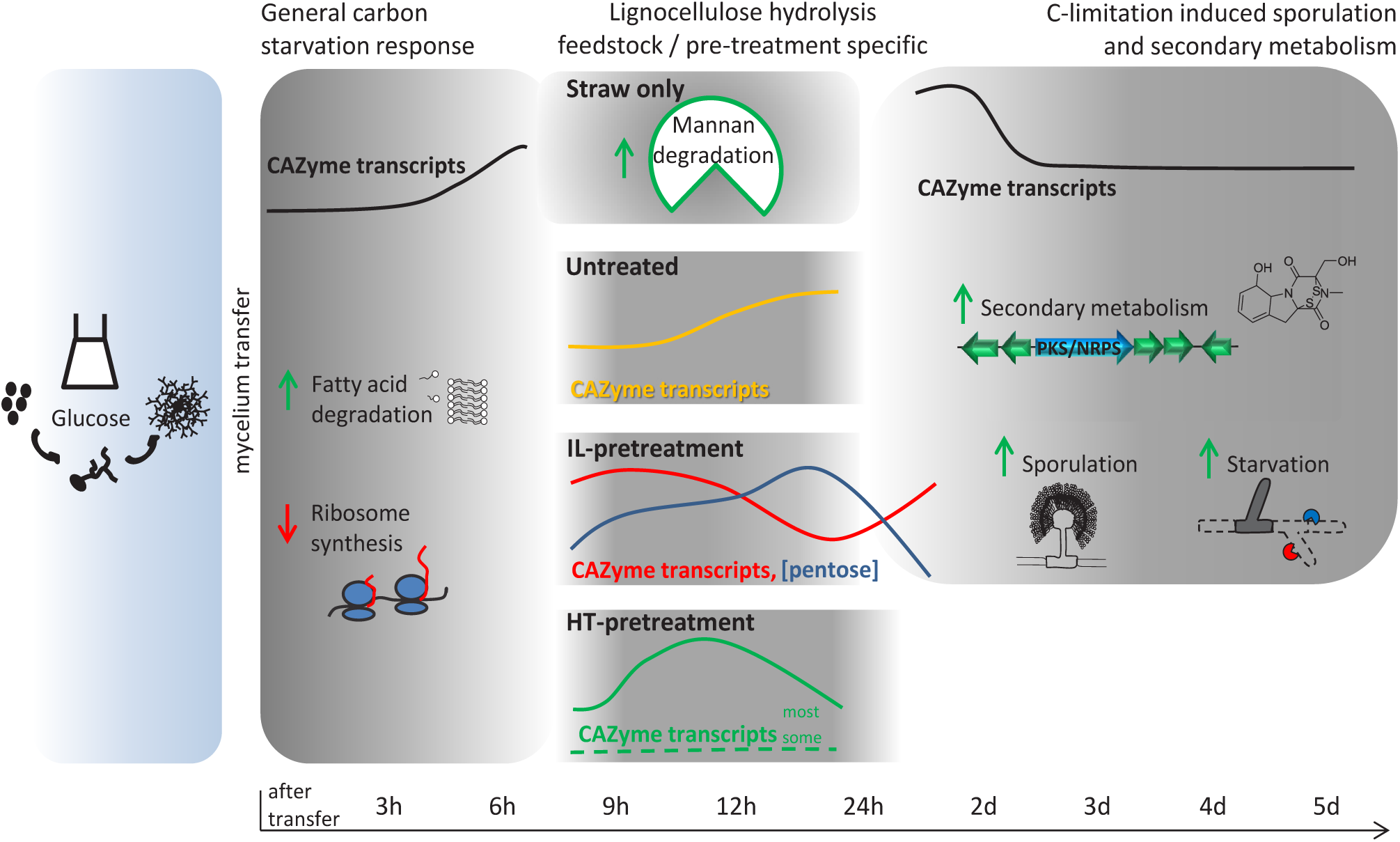
Illustration of physiological stages of the *A. niger* response to (pretreated) substrates including both conserved and substrate-specific responses. After transfer to lignocellulose, *A. niger* reacted with a general carbon starvation response, followed by pretreatment and feedstock specific elements such as mannan degradation on straw but not *Miscanthus*, followed by a general carbon-limitation induced response with sporulation and induction of secondary metabolite clusters. Stand-out features of the response are indicated for each condition. Green and red arrows represent processes that were increased or decreased, respectively. CAZy transcript levels and pentose concentration are represented as average (for all or stated conditions) percentage of the maximum with identical y-axes in all instances. The width of blue and grey panels represent the covered time-period, marked on the time-scale as h after transfer to lignocellulose.

Indications for starvation and stress in the late time point cultures were found through enrichment of the following GO-terms among genes with increased transcript levels: retrograde vesicle recycling in the Golgi, (non-glycolytic) fermentation and ionic osmotic (saline) stress responses, cell death, DNA modification, lipid metabolism and fungal cell wall acting enzymes (Fig. 6b). Starvation and stress occurred later in the IL pretreated compared to HT or untreated cultures, consistent with ILS and ILM having increased digestibility. Fungal cell wall-acting CAZymes encoding genes that are responsive to carbon starvation, such as *cfcA/chiB* and *engA* have high transcript levels from 12 h (HT, KMM), 24 h (KMS) or 2 d (IL) (Fig. 6c) with corresponding accumulation of proteins (Fig. 6d). Starting from 24 h (HT, KM) or 2 d (IL) cultures, transcript levels were strongly increased for genes encoding sporulation transcriptional activators (*brlA, abaA*), pigment synthetic enzymes (*fwnA, brnA, olvA*), hydrophobins (predominantly *hfbA, hfbB, hyp1*) and sporulation-specific cell wall modifying CAZymes (*ctcB, bgtD*), indicating sporulation was taking place (Fig. 6e). GO-terms for breakdown of aromatic amino acids (phenylalanine, tyrosine, tryptophan) were enriched at most of the time-points without high monomeric sugar levels. These GOterms share common genes that belong to the tyrosine-phenylalanine metabolism gene cluster. Transcription of these genes was co-regulated and was most abundant on later (≥ 24 h) time points. Together this indicates that the late time points represent carbon-limited fungal cultures with limited growth, which activated survival strategies such as sporulation and secondary metabolism.

### Novel and known secondary metabolite gene clusters induced at later time points

Fungal secondary metabolites attract high interest due to their potential bioactive properties (65). *A. niger* contains many secondary metabolite gene clusters (65,66) of which for many their products are so far unknown. The production of secondary metabolites has been reported in some instances to be linked to starvation and sporulation but for a large number of metabolite clusters no conditions have been identified under which they are expressed. Our lignocellulose time-courses represent a range of novel conditions that could stimulate the fungus to generate secondary metabolites and we therefore assessed expression of secondary metabolite generating gene clusters predicted in the *A. niger* genome.

The GO-term for secondary metabolism was enriched most notably in the ILM and ILS time courses (Fig. 6b). The gene cluster encoding proteins for production of the toxin fumonisin (An01g06810-An01g06970) was strongly induced on the 2 d ILS cultures and to a more moderate level in 2 d ILM. Gene cluster An09g00520-An09g00620, containing a gene with strong similarity to *gliC* of the *A. fumigatus* gliotoxin synthesis gene cluster (67), showed a pronounced increase in transcription levels at the 9 h KMS and ILM 2 d time points, as well as some increases on ILS 2 d (Fig. 6f). This peculiar transcript profile was shared with two genes (An01g06150 and An08g06810) involved in the shikimate pathway, which generates chorismate that feeds into biosynthesis of aromatic amino acids such as phenylalanine, one of the two amino acids that are condensed to produce gliotoxin. Metabolic mapping also pointed towards enrichment of the shikimate pathway (part of KEGG pathway rn00400). The secondary metabolite cluster An11g06420-An11g06490 showed strong induction from IL 2 d and 3 d, as well as some expression on later IL time points and on KMS from 24 h onwards. This cluster was also induced in *A. niger* in mixed-species cultures on wheat straw (68). An orthologous hybrid PKS/NRPS is required for production of pseurotin A in *A. fumigatus*, and is induced in response to hypoxia and the anti-fungal drug voriconazole (66,69,70). Expression of the uncharacterised cluster An12g07060-An12g07110 (71) increased to considerable levels on late cultivation time points (Fig. 6f). Overall, the transcriptomic data indicated activation of the production of potentially new as well as known secondary metabolites, confirming that cultivation with plant biomass can provide unique conditions for the induction of secondary metabolite gene clusters in fungi.

## Conclusions

The transcriptional response of *A. niger* to lignocellulose adapted over the cultivation time-course, and can overall be divided into three stages: an initial (3 - 6 h) conserved response, followed by a substrate and pre-treatment-specific response (generally 9 – 24 h) and a final largely conserved response (generally 24 h - 5 d) albeit with variable elements, as summarised in (Fig. 3).

Initially fatty-acid β-oxidation was activated and a role in timely establishment of lignocellulose degradative capacity was identified. This was followed by activation of a substrate hydrolytic stage that was mainly directed by substrate properties such as composition and digestibility. Subsequently, sporulation was activated, and the transcriptome data indicated this was in response to reduced metabolic activity and carbon limitation in the culture. This study emphasises the value of high-resolution time-courses when studying fungal responses to lignocellulose, to ensure confidence in identification of all involved enzymes that may be expressed, and importantly, to understand more broadly the range and dynamics of the fungal physiology on a systems level. Ultimately, improved understanding of these responses should improve the efficiency of strain and enzyme developments for renewables-based biotechnology.

We identified a range of secondary metabolite gene clusters that were activated during exposure to lignocellulose, including multiple clusters so far without product characterisation. Secondary metabolite gene clusters attract significant interest for generating novel potentially bioactive compounds and as a source of bio-catalytic enzymes. We are therefore confident that the expression conditions for these novel clusters identified here will receive interest by providing opportunities for their further characterisation and exploitation.

## Methods

### Substrates and pretreatments

*Miscanthus* (cv. Giganteus) was ground to pass an 0.5 mm sieve as described previously (31). The ionic liquid (IL) pretreatment was performed using 1-ethyl-3-methylimidazolium acetate ([C2mim][OAc]) for 2 h at 160 °C and hydrothermal treatment was performed at 200 °C for 2 h, as described in detail previously for wheat straw (31). The compositional and structural analysis of substrates was performed as described in detail previously for wheat straw (31), employing total acid hydrolysis assay followed by analysis of soluble sugar monomers in the hydrolysate by high-performance anion exchange chromatography with pulsed amperometric detection (HPAEC-PAD), and wide-angle X-ray diffraction. Digestibility of the substrates was assessed in saccharification assays with a commercial enzyme mixture (Celluclast (Sigma) and Novozyme 188 (Sigma), 1:1 v/v), as well as by measurement of residual xylose in solids of shake-flask cultures during fungal exposure (31).

### Fungal cultivation conditions

Cultivations were performed as described before (31). Briefly, 1 % w/v glucose cultures were inoculated with *A. niger* N402 spores at a final concentration of 1×10^6^ ml^−1^and incubated at 28°C for 48 h. The mycelia were collected and 1.5 g wet weight was transferred to flasks containing one of the following carbon sources at a 1 % w/v final concentration in AMM: glucose, untreated *Miscanthus*, ionic liquid pretreated *Miscanthus* or hydrothermally pretreated *Miscanthus*. The cultures were incubated for 3 h, 6 h, 9 h, 12 h, 24 h, 2 d, 3 d, 4 d or 5 d. Mycelia were collected for RNA extraction and the filtrate of the culture was flash frozen in liquid nitrogen and stored at -80 °C for subsequent sugar and targeted proteomics analyses, both performed as described (31).

### Gene expression analysis

RNA extraction, Illumina paired-end sequencing, read mapping to the *A. niger* ATCC 1015 strain and statistical analysis were all performed as described in detail before using a DESeq2 *p*_adj_ value < 0.05 to identify differentially expressed genes between pairs of conditions (31). The obtained data were analysed together with those obtained from cultivation with untreated and pretreated wheat straw (31) under identical conditions. Table S6 contains the FPKM values for each sample with *Miscanthus* as feedstock as well as the DESeq2 analysis for the comparison of each condition with the Glu 48 h control. Clustering and Gene Onotology (GO) enrichment analysis was performed as described before (31) using the combined dataset including wheat straw and *Miscanthus* derived samples. In short, hierarchical clustering of log transformed gene expression values (FPKM) of all 47 conditions with R package gplots. GO enrichment analysis was performed using the Bioconductor package GOseq (v1.18.0) (72) using GO annotations from the FetGOat resource (http://www.broadinstitute.org/fetgoat/) (73), and comparing to the Glu 48 h control for each condition the genes with a log2 fold change (i.e. DESeq2 of £ -1 or ≥1), *p*_adj_ £ 0.05 and an FPKM ≥ 1 in at least 1 condition. PCA analysis was performed using the MetaboAnalyst package (74,75), using log transformed, mean centered FPKM values of triplicate conditions. Genes with high calculated loadings are analysed for GO-enrichment using g:Profiler (76), results were summarised using REVIGO (77) and subsequently visualised using https://www.jasondavies.com/wordcloud/ to generate word clouds.

### Mapping transcript level data to a genome-wide metabolic model

In order to analyse gene expression data on a metabolic network we used a CoReCo created genome wide metabolic model of *A. niger* (78). For each enzyme commission (EC) number in the model an arbitrary enzyme expression value was created by taking the expression value of the responsible gene or taking the highest value when multiple genes were mapped to one EC number. The resulting EC expression data matrix was then explored using principal component analysis and Bayesian Hierarchical Clustering (79), followed by mapping of the EC cluster content to pre-defined pathways in KEGG (80) and Metacyc (81) databases. Enrichment of EC cluster members on predefined pathways was quantified with the hypergeometric test.

### Analysis of gene expression in *farA* deletion strain

The wild-type strain N402 and two independent *farA* deletion strains (MA537.3 and MA537.4, kindly provided by Dr A. Ram, University of Leiden, the Netherlands) were grown in liquid cultures in triplicate as described above using 1 % ball milled wheat straw as substrate. After 9h and 24 h, mycelium was harvested and RNA was isolated as described above. qRT-PCR was performed as described previously (31) but with PCR products as a standard, using 0.5 µl of cDNA and the primers specified in Table S7. Gene expression values are given as % of expression of *actA* and *sarA*. Statistically significance of differences in gene expression was tested with a one-way ANOVA with post-hoc multiple comparisons test.

### Enzyme assays

For the enzyme assays with AZCL dyed galactomannan (I-AZGMA, Megazyme), appropriate volumes of the concentrated culture filtrates were incubated for 3 h at 28 °C with 150 rpm shaking with 1 % w/v concentration of the dyed substrate in 100 mM sodium acetate pH 4.5 in a total volume of 250 μl in 96 well plates. Reactions were stopped with 2% Trizma base as per manufacturer’s instructions and absorbance was measured at 590 nm with a plate reader.

## Abbreviations

CAZy: carbohydrate-active enzymes
CCR: carbon catabolite repression
FPKM: fragments per kilobase of gene model per million mapped fragments
GO: gene ontology
HPAEC-PAD: high-performance anion exchange chromatography with pulsed amperometric detection
HT: hydrothermal
HTS: hydrothermally pretreated straw
HTSM: hydrothermally pretreated *Miscanthus*
IL: ionic liquid
ILS: ionic liquid-pretreated straw
ILM: ionic liquid-pretreated *Miscanthus*
KMS: knife-milled (untreated) straw
KMM: knife-milled (untreated) *Miscanthus*
MWCO: molecular weight cut-off
NRPS: Nonribosomal peptide synthetases
p_adj_: *p* value from DESeq statistical analysis adjusted for multiple hypothesis testing
PCA: Principal Component Analysis
PKS: polyketide synthases
SignalP: signal peptide
TCA: tricarboxylic acid

## Declarations

### Availability of data and materials

The RNA-seq reads supporting the conclusions of this article are available in the BioProject repository accession PRJNA250529, http://www.ncbi.nlm.nih.gov/bio-project/PRJNA250529. Other datasets supporting the conclusions of this article are included within the article and its additional files.

## Authors’ contributions

STP, SD, DBA, GAT and BAS conceived the RNAseq experiment. PD, JMvM, MK performed fungal culture work and RNA and protein sample preparation. RI and SG prepared lignocellulosic substrates and performed compositional analyses. IVG, KWB, EL and CYN co-ordinated the RNAseq work at the JGI. AL and VRS performed bioinformatic analysis at the JGI. MJB performed clustering, enrichment analysis using the RNA-seq data. JMvM performed multivariate analysis of RNAseq data. PD and JMvM analyzed the data. PD, JMvM, DBA wrote the manuscript. RR, PD and JMvM performed experiments with the *farA* deletion strain. CJP coordinated the targeted proteomics work at JBEI. CJP and LC performed the targeted proteomics work at JBEI. MA performed mapping of transcript data to the metabolic model.

## Acknowledgements

We thank Dr. Sarah Purdy for the *Miscanthus* substrate and Yvonne Arendsen, Dr. Noel van Peij and Dr. Herman Pel (DSM, the Netherlands), Mark Arentshorst and Dr. Arthur Ram (Leiden University) for making the *farA* deletion strains available.

## Supplementary Data

**Figure S1. Characterisation of untreated and pretreated *Miscanthus* substrates. a** Carbohydrate monomer composition of substrates, **b** substrate crystallinity with the predominant type of cellulose indicated **c** saccharification of substrates using commercial enzymes given as mean (n=3) ± SE of released glucose and xylose after 72h, **d** xylan in initial substrates and residual xylan from the solids recovered from the fungal cultures after 24 h and 5 d (where available), given as mean (n=3) ± SE of biological replicate flasks or for the initial substrates as the mean ± SE of 2 technical hydrolysis replicates, **e** monosaccharide concentration in the fungal culture filtrates over time, given as mean (n=3) ± SE with A/C only = autoclaved substrate only.

**Figure S2. Expression of CAZyme encoding genes over *Miscanthus* time courses. a** The total number of genes encoding plant-polysaccharide active CAZymes and that are significantly induced in cultures with untreated and pretreated *Miscanthus* as compared to the Glu 48 h control (DESeq *p p*_adj_ < 0.05, FPKM of ≥ 50 on lignocellulose and log2 FC of ≥ 3). Values are mean **±** standard errors (n=3). **b** The proportion of transcripts from these CAZyme genes as percentage of total FPKMs. Error bars represent standard errors (n=3). Colors indicate different culture condition groups and values in light grey are a comparison to the same data when using wheat straw as feedstock (31).

**Figure S3. Targeted proteomics of selected proteins over the Miscanthus time courses.** Relative abundance of the indicated enzymes are given as peak area corrected for total protein amount in culture filtrate. Values are given as mean ± st. error (n=3) and gene identifiers corresponding to the proteins are given in Table S5.

**Table S1. File with lists of enriched GO terms in genes with reduced transcript abundance.** Enriched GO terms (and the genes in the list responsible for the enrichment) from the enrichment analysis of the genes that were repressed in the comparison of the Glu 48 h control with untreated and pretreated *Miscanthus*.

**Table S2. File with lists of enriched GO terms in genes with increased transcript abundance.** Enriched GO terms (and the genes in the list responsible for the enrichment) from the enrichment analysis of the genes that were induced in the comparison of the Glu 48 h control with untreated and pretreated *Miscanthus*.

**Table S3. Mapping transcript level data to a genome-wide metabolic model.** Gene transcript levels were mapped to enzyme activities, and converted to an expression value per enzyme activity (ECx.x.x.x) number. Genes mapped to the EC number are indicated. Pathways are listed that were found significantly enriched in clusters after mapping of the activities to databases with metabolic pathways, as well as an overview of all KEGG pathways to which EC numbers were mapped.

**Table S4. PCA loadings and enrichment of GO-terms in genes distinguishing sample clusters.** Loadings of genes on the first 10 components are given of the PCA analysis of all cultivation conditions, and of those from the 9h and 12h time points of the Miscanthus time course. Results are included of GO-enrichment analysis (via gProfiler) of genes with high loadings on the PCA components corresponding to the sample clusters identified in Fig2, i.e. Cluster 1 PCA1<0 PCA2<0; Cluster 2 PCA1>0; Cluster 3 PCA1<0 PCA2>0; Cluster 3a PCA1<0 PCA>0 PCA3>0. GO-enrichment analysis are also included for 9h and 12h Miscanthus conditions PCA1<0 PCA2<0 (HTM specific) and for PCA4>0 (Wheat straw specific).

**Table S5. Data supporting CAZyme expression analysis.** Gene lists used to generate Venn diagrams depicted in Fig5c and d are given including Venn diagram intersections, the CAZyme encoding genes, FPKM transcription levels and classification underlying Fig. 4e are given, as are gene identifiers corresponding to proteins targeted with proteomics (FigS3).

**Table S6. Gene expression as identified by RNA-Seq, and corresponding statistical analysis.** Gene identifiers and annotations are provided along with the FPKM values for each sample and condition, the fold change and *p p*_adj_ values as calculated by DESeq2 for the comparison between the Glu 48 h control and the other conditions.

**Table S7. Primers used for qRT-PCR analysis.** Nucleotide sequences of primers are given together with gene identifiers

