## Supplementary figures and images for "Succession of physiological stages hallmarks the transcriptomic response of fungus *Aspergillus niger* to lignocellulose"

### Supplemental Figure 1

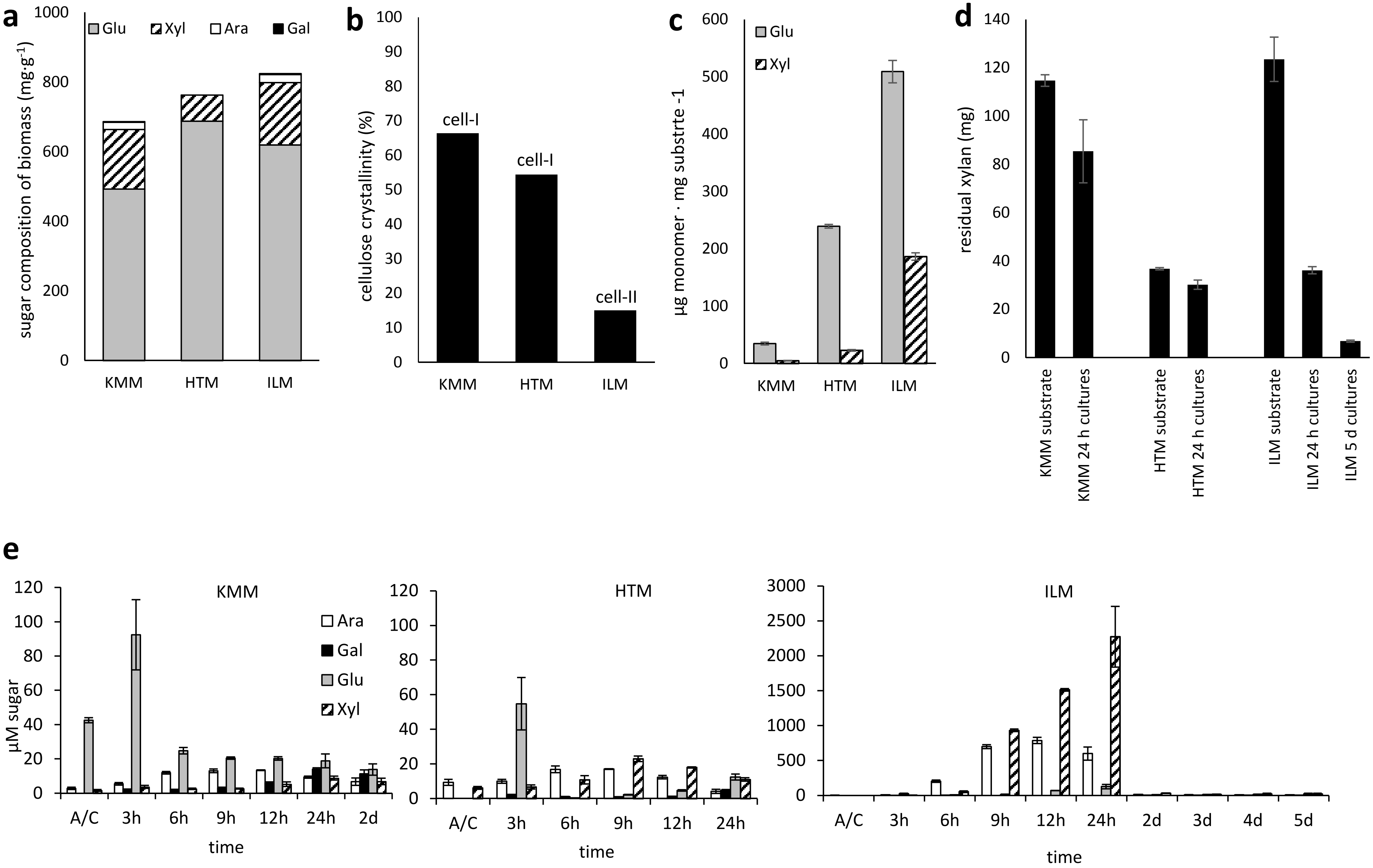

### Supplemental Figure 2

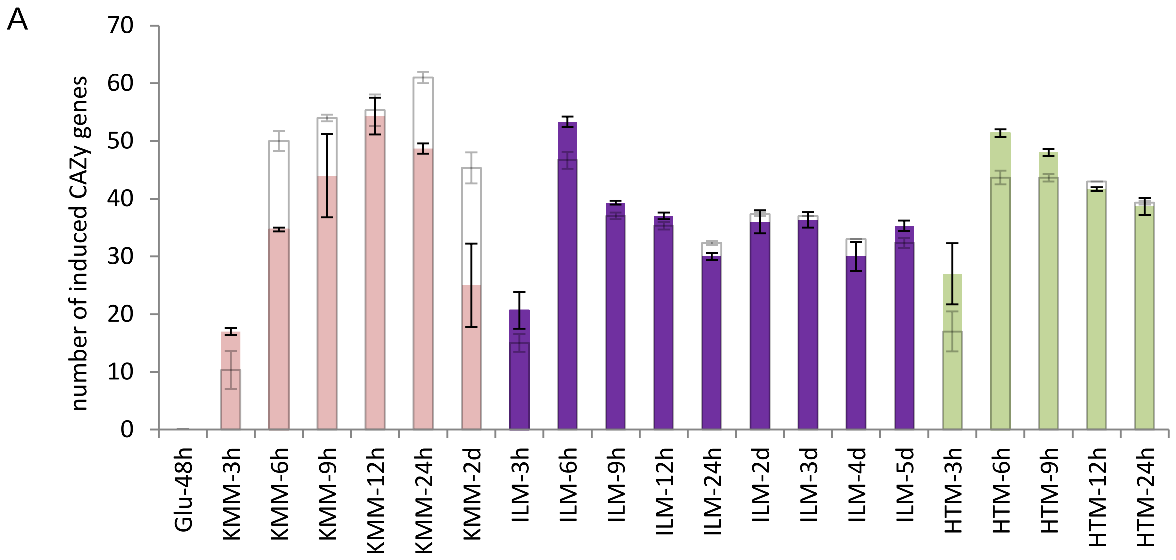

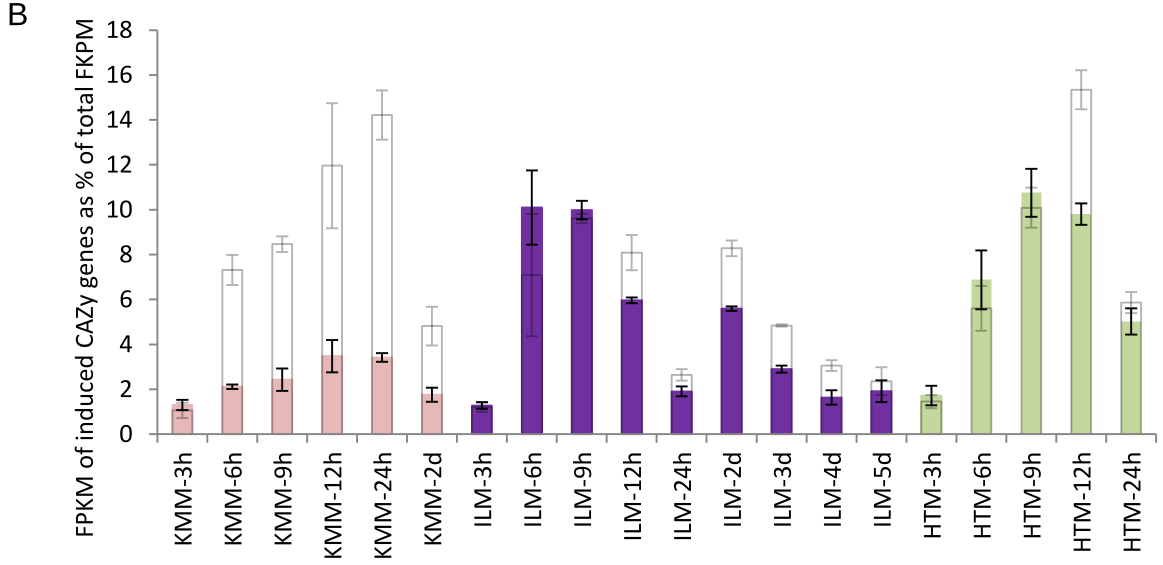

### Supplemental Figure 3

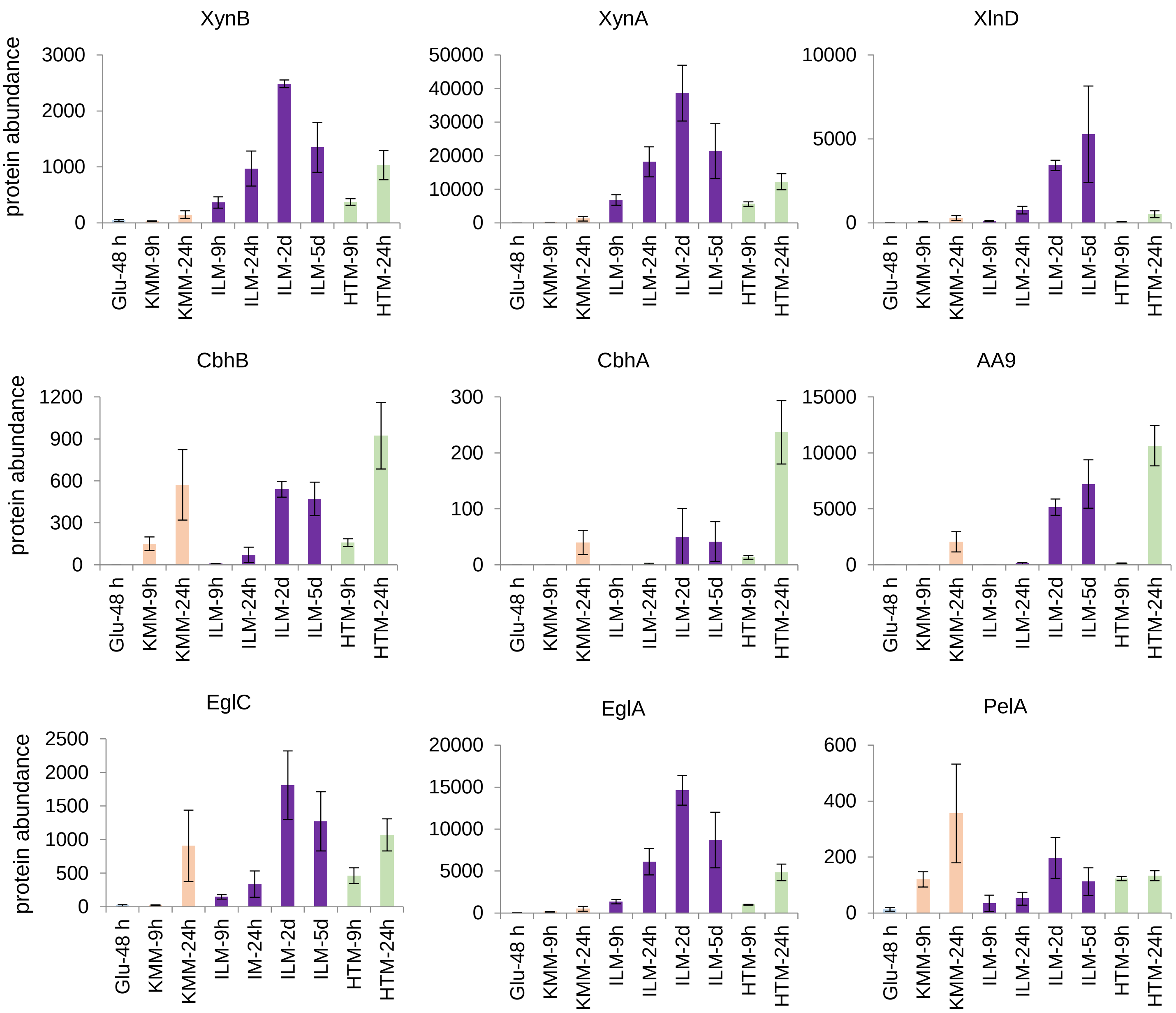

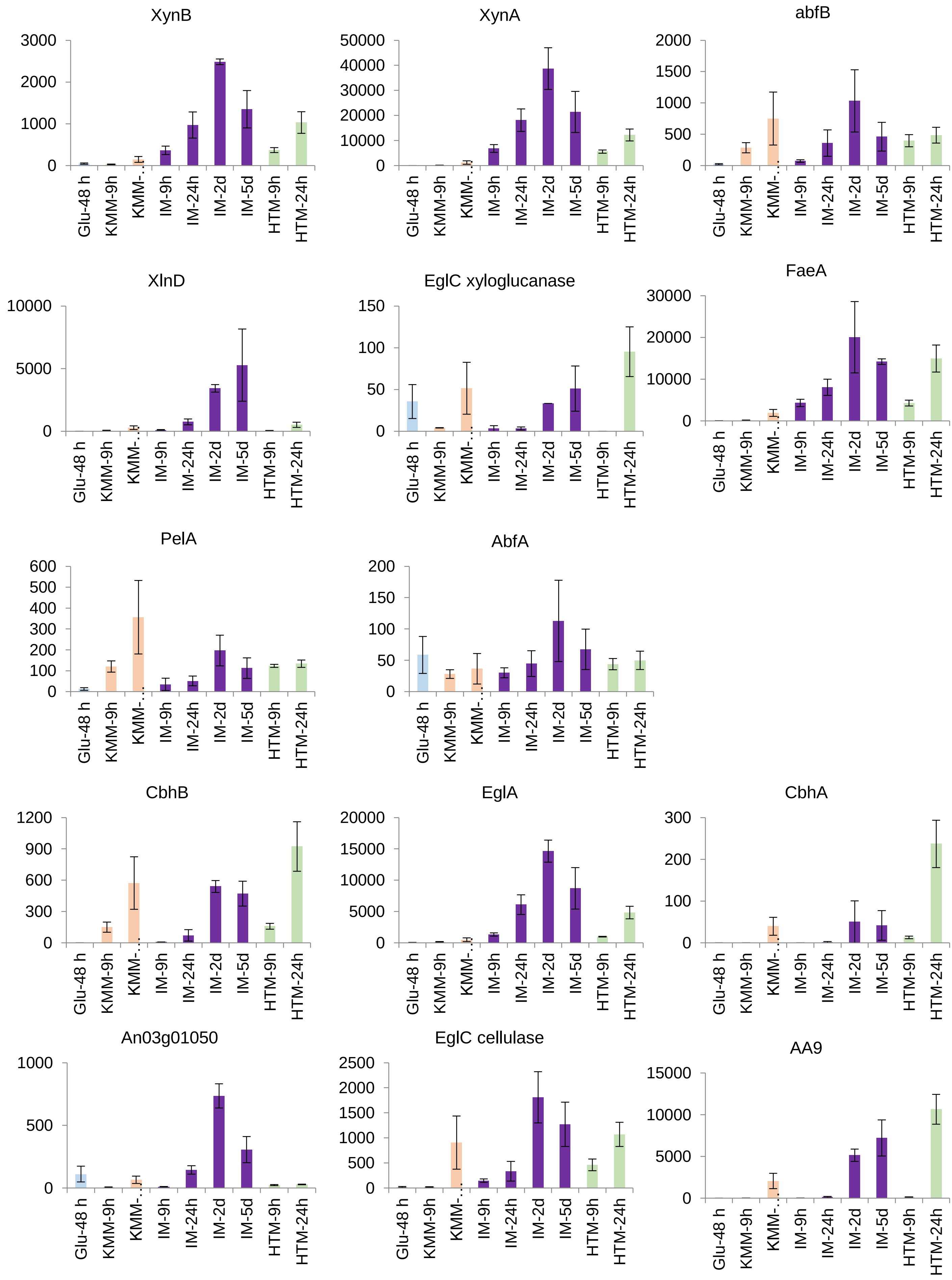

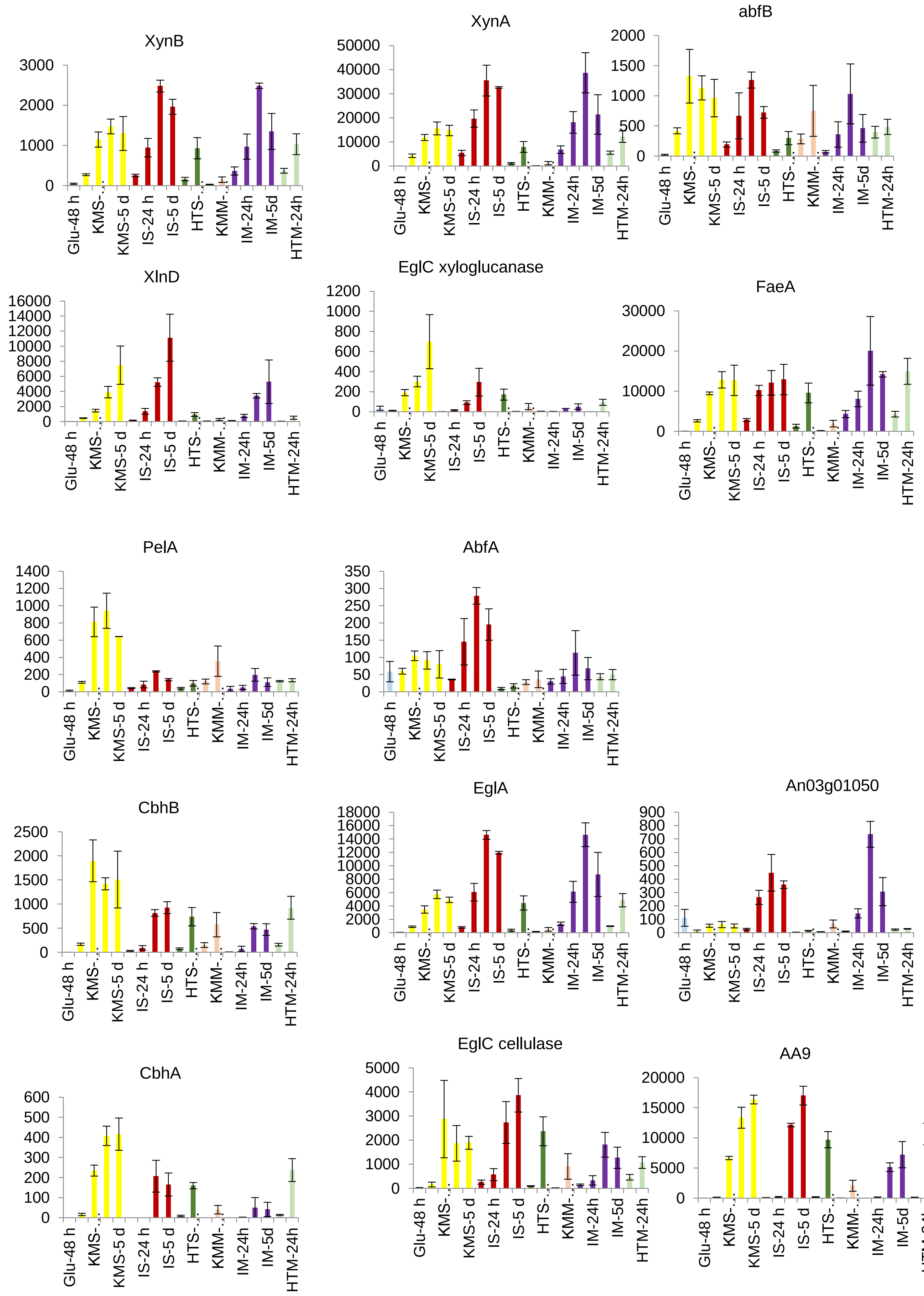
